# Gene target discovery with network analysis in *Toxoplasma gondii*

**DOI:** 10.1101/397398

**Authors:** Andres M. Alonso, Maria M. Corvi, Luis Diambra

## Abstract

Infectious diseases are of great relevance for global health, but needed drugs and vaccines have not been developed yet or are not effective in many cases. In fact, traditional scientific approaches with intense focus on individual genes or proteins have not been successful in providing new treatments. Hence, innovations in technology and computational methods provide new tools to further understand complex biological systems such as pathogens biology. In this work, we propose a system biology approach to analyze transcriptomic data of the parasite *Toxoplasma gondii*. By means of an optimization procedure, the phenotypic transitions between the stages associated with the life cycle of the parasite were embedded into the dynamics of a gene network model. Thus, through this methodology we were able to reconstruct a gene regulatory network able to emulate the life cycle of the pathogen. The community network analysis has revealed that nodes of the network can be organized in seven communities which allow us to assign putative functions to 339 previously uncharacterized genes, 25 of which are predicted as new pathogenic factors. Furthermore, we identified a small subnetwork module that controls the parasite’s life cycle. These new findings can contribute to understand of parasite pathogenesis.

## Introduction

Toxoplasmosis is a zoonotic disease that affect almost one third of the global population^1^. This condition is caused by the obligate intracellular parasite *Toxoplasma gondii*, which transits a complex life cycle. It develops an asexual phase in mammals and birds where the parasite cell can adopt an invasive and rapidly dividing form, the tachyzoite, and a latent form which is encysted in the host, the bradyzoite^2^. When bradyzoites are ingested by members of the *felidae* family, the definitive host, they differentiate into merozoite -an invasive and asexual form that will originate sexual gametocytes- and finally a sporulated form in oocysts, the sporozoite, as illustrated in Figure 1A. Thus, the transitions between the different stages of the cycle allow the pathogen to adapt to diverse contexts by modulating its virulence ans pathogenic potential^3^. While the stages of the biological cycle of *T. gondii* are characterized, the mechanisms that regulate the transitions between them are not completely understood. Different studies were directed to understand the phenomenon postulating that epigenetic regulation, changes in gene expression and subsequent activation/deactivation of genetic networks play a relevant role in the conversion from one stage to another^4^.

**Figure 1.**
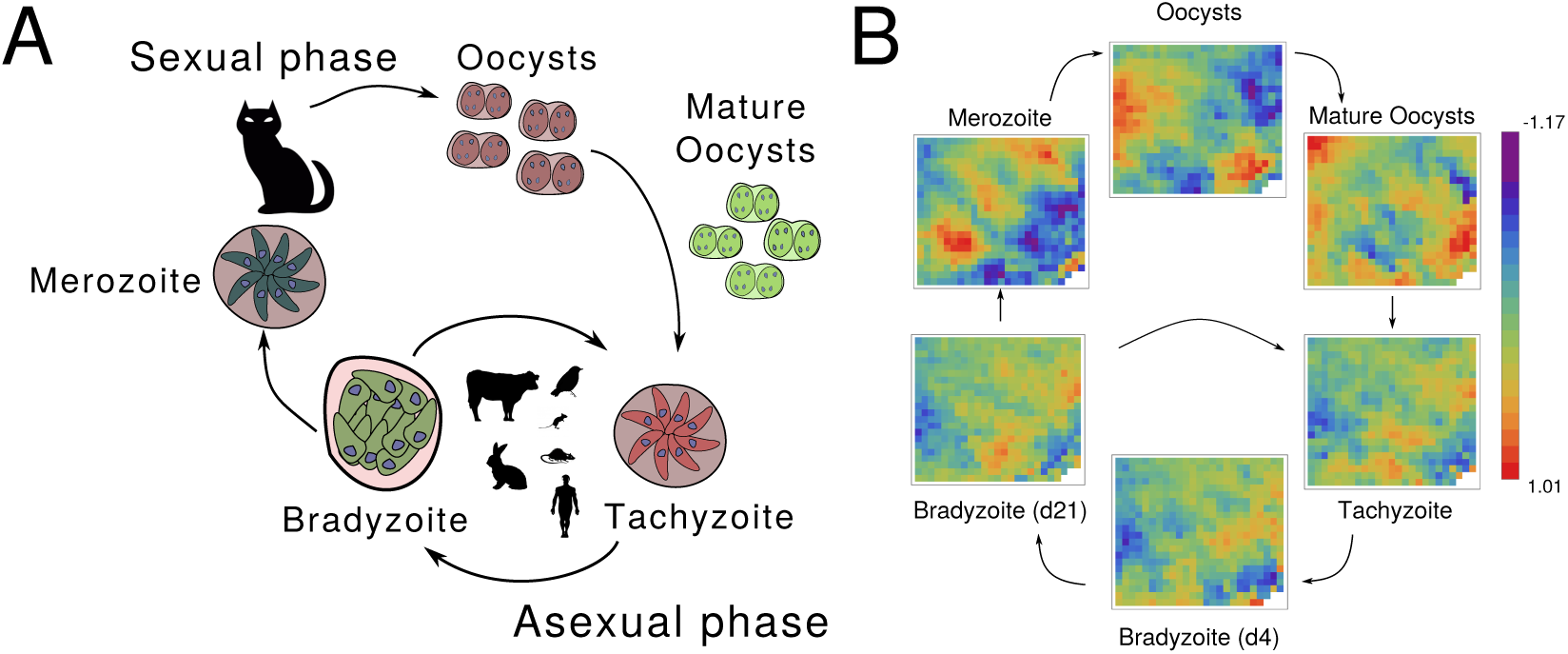
*Toxoplasma gondii* life’s cycle. (A) A schematic representation of the parasite biological cycle; (B) Transcriptional snapshots of the parasite’s cycle (five stages). After a dimensional reduction procedure, we have found that microarray dataset can be represented by 545 variables. Each of these variables (cells in the 22*×* 25 array) corresponds to the intra-cluster average of the log-transformed relative expression level of the genes that belong to the corresponding cluster. Since gene assignment to clusters is the same for all states, the arrays can be directly compared between them.

In order to understand how the *Toxoplasma* cycle is orchestrated, several systematic approaches have been implemented which are based on the application of high-throughput technologies (HTTs) in the field of epigenetics, genomics and proteomics. The protocols used include Chromatin immunoprecipitation (ChIP) in conjunction with microarray technologies (ChIP-chip)^5, 6^, high-throughput sequencing (ChIP-seq) and gene expression studies based on microarray or sequencing technologies (RNA-seq)^7, 8^. Given the range of experimental conditions and the typical performance of such high performance techniques, a new challenge arises: organize and analyze resulting information from new technologies in a coherent work framework. The methodologies mentioned above provide almost complete observations of complex biological systems and can lead to a deeper understanding of the problem at the systems level. Consequently, understanding biological systems requires HTTs data products integration which are used to build quantitative models. Systems biology is an emergent and multidisciplinary field that proposes new and rational approaches for the analysis of HTT-derived information in the field of infectious diseases^9^. One of these goals involve the inference of gene regulation networks (GRNs) from large amounts of information, since it allows modeling the dynamics of complex systems in a single conceptual framework^10–12^. GRNs are dynamic systems whose states are determined by the expression levels of each gene or groups of genes (nodes), while the edges, or links, between nodes represent regulatory interactions; the structure of the network is then defined as a graph^13^. Once the network is reconstructed it is possible to address a number of different biological and biomedical questions such as the dissection of a key gene circuit involved in cellular differentiation^14^, the study of phenotypes related to health conditions, the development of new therapies, the design of perturbation experiments^15^ and interpretation of direct gene interactions such as transcription gene regulation^16,17^. However, uncovering the GRN architecture represents a very difficult task, due to the large number of genes and the limited amount of data available, but also by the nonlinear dynamics of regulations, the noisy readouts of expression levels, and other factors that are part of the challenge^18^. In this sense, a computational technique that allows the reconstruction a GRN from gene expression levels that overcome several major obstacles has been recently developed^12^ and applied to *T. cruzi*.

Here we apply a GRN approach to study the life cycle of *Toxoplasma gondii*, by integrating transcript expression data from sexual and asexual phenotypes Figure 1B. Our proposed framework helps to uncovered the underlying architecture of the network that supports the steady states associated with six phenotypic stages of *T. gondii* and the transitions between the parasite’s life cycle stages in response to environmental cues. The method was efficient to elucidate master key regulators involved in the analyzed phenotypical transitions. Most of the genes that are part of the subnetwork have not been characterized yet, while the presence of four dense granule proteins (GRA1, GRA2, GRA6, and GRA12) is highlighted. Finally, *in silico* perturbation experiments propose these key genes for future experimental studies in the tachyzoite to bradyzoite differentiation. Furthermore, by combining clustering methods and communities analysis it is possible to infer biological processes associated to these uncharacterized genes. While genes that are co-expressed tend to take part in the same processes and perform similar or complementary functions^18^, the inference of communities in the network allows to predict putative functions within the network.

We believe that the study of pathogen’s life cycles by gene network models leads to a thorough understanding the understanding of signaling pathways and their actors, being a powerful predictive tool for new molecular targets and diagnosis development as well as to assign functions to uncharacterized genes.

## Results

### Modeling the gene regulatory network of *T. gondii*

In order to model the *T. gondii* GRN we assumed that the system’s state was represented by the vector *x*(*t*) corresponding to the expression levels of *N* gene clusters measured at time *t*. The GRN dynamic was modeled by a first order Markov model, where future states were linearly dependent on the present state, following this equation:

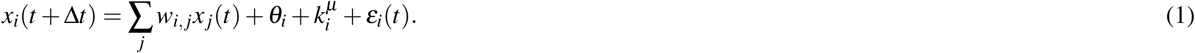

Thus, the evolution of the system is governed by the matrix **W** and the external perturbations by **k**^*µ*^. The elements of the weighted connectivity matrix, *w*_*i,* *j*_ indicate the type and strength of the influence of gene *j* on gene *i* (*w*_*i* *j*_ *>* 0 indicates activation, *w*_*i* *j*_ *<* 0 indicates repression, while 0 indicates absence of regulation). The influence of environmental cues on genes are represented by 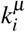. The next step consists of determining which nodes are affected by the environmental cues, and how they are affected. To this end, not only did we consider the expression-profile data set, but also some *a priori* known biological facts: (i) the parasite’s life cycle has five stages: immature oocysts, mature oocysts, tachyzoite, bradyzoite and merozoite; (ii) there are four possible transitions between these five stages, promoted by different environmental cues; (iii) each steady state exhibits some level of fluctuations around it; and (iv) the connectivity matrix is a sparse matrix. The available transcriptome data consist in six data points over the life cycle of *T. gondii*: oocysts day 0 (Od0), oocysts day 10 (Od10), tachyzoite day 2 (Tzd2), bradyzoite day 4 (Bzd4), bradyzoite day 21 (Bzd21), and merozoite of cat #52 (Mc52). Notice that the four external differentiation signals considered here, **k**^*µ*^ with *µ* = 1, 2, 3, 4, are associated to the following phenotypic transitions: Od0 → Od10, Od10 → Tzd2, transition trajectory Tzd2 → Bzd21. Following Bzd21 and Bzd21→ Mc52, respectively. The data point Bzd4 is part of the transition trajectory Tzd2 → Bzd21. Following^12^, we implemented a two-step reverse engineering protocol, focusing first on embedding the six data points into the dynamics of the model, regardless of the transitions between these states. In a second step, we concentrated on uncovering the environmental cue effectors considering the transitions between the parasite’s life cycle stages, while using a connectivity matrix consistent with steady states.

### Embedding the steady states

In order to embed the basins of attraction which correspond to each of the parasite’s stages, we have considered the model described in Eq.1 without external perturbations, and applied the singular value decomposition (SVD) procedure to compute connectivity matrix **W**, as indicated in the corresponding Methods section. The elements of matrix **W** are continuous variables, though many of the inferred matrix elements are not much value different from zero. Consequently, the number of predicted edges of the GRN is quite high compared to the number of edges in known regulatory networks^19, 20^. With the aim to obtain a sparse version of the connectivity matrix, i.e. a **W** matrix whose elements should be either 0 or another reliable value different from 0, we considered a kind of bootstrap method. In this sense, we constructed an ensemble of 500 training sets by adding different noise realizations to the different stages. By computing the minimum *L*_2_ norm solution for each training set, we obtained a probability distribution for each weight, *P*(*w*_*i,* *j*_). In this manner, we were able assess the significance of the weighted values by performing a location test to prune the non-significant weights, and constructing a sparse connectivity matrix, **W**_*ss*_, which supports the data set. At the significance level of 0.01 we found 17, 873 links between genes, i.e. around the 94% of the elements of **W**_*ss*_ were null. To quantify how the dynamics of the GRN, with the matrix **W**_*ss*_, was able to capture the parasite’s stages as basins of attractions of the system, we computed the overlap between the vectors that represented the actual state of the network and the target stage. Mathematically, the overlap between vectors *x* and *y* is defined as *x · y/| x | |y |*. Figure 2A depicts the trajectories that illustrate the dynamics of our model in the space spanned by the three principal components. A zoomed view over the stages Tzd2 allows to appreciate that the trajectories (black lines) fluctuate around the corresponding stage represented by a colored circle (Fig. 2B). This means that whenever the system is in a given basin, it will remain inside that basin as long as there are no external perturbations. For each trajectory we computed the overlap between the vectors corresponding to the state of the system at time *t*, and the ones corresponding to each target stage. Figure 2C depicts the time course of such overlaps illustrating, in a more quantitative manner, how the trajectories fluctuate around each parasite’s stage.

**Figure 2.**
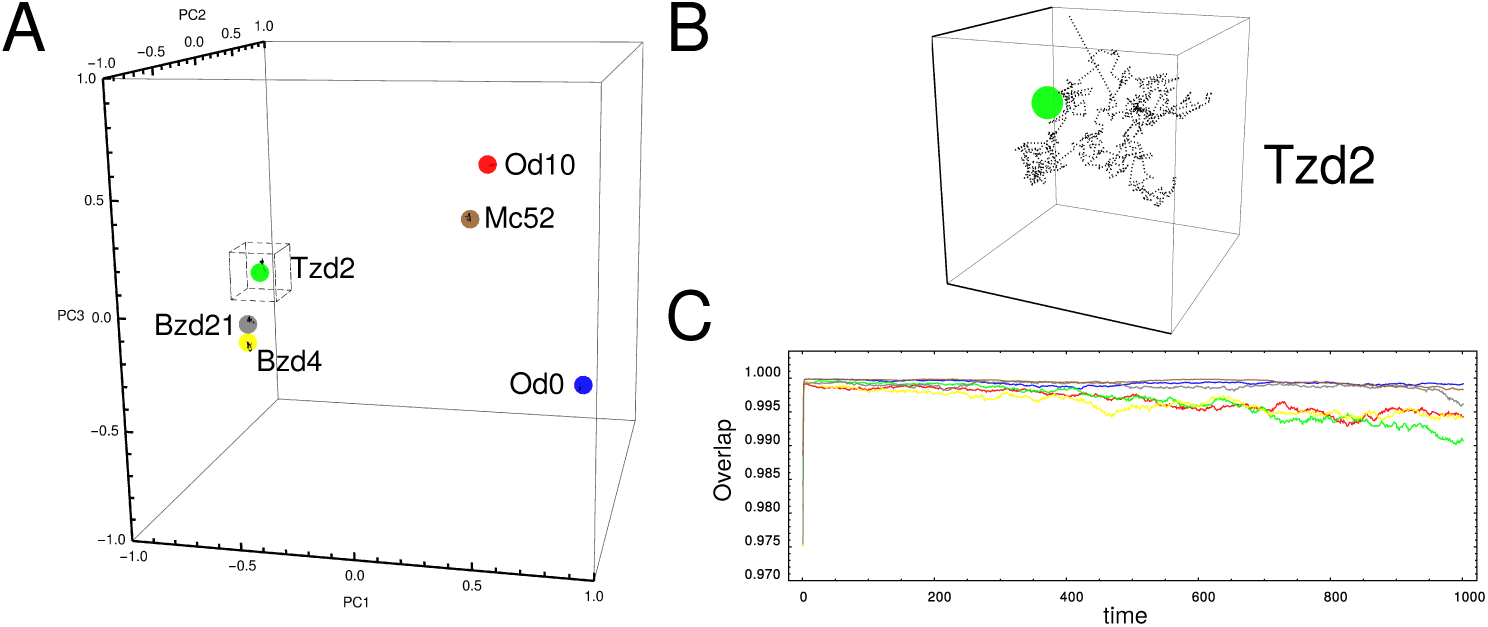
Stability of the steady states. (A) The plot shows the positions of the steady states of the parasite’s life cycle in the space spanned by the three principal components. The black trajectories around each stage are the result of simulations conducted using the model Eq.(1) without external cues. In each case a slightly perturbed steady state was used as the initial condition; (B) Temporal behavior of the overlap between the state of the system at time *t* and the target steady state: Od0 (blue), Od10 (red), Tzd2 (green), Bzd4 (yellow), Bzd21 (gray) and Mc52 (brown); (C) A zoom view over the trajectory around the Tzd2 steady state. One can observe that the system fluctuates around the corresponding this state.

The visualization of the obtained network is a challenging task, even with the low average node degree associated to our sparse GRN (*∼*6%). In order to overcome this obstacle, we displayed only a small fraction of the nodes grouped in seven communities. Figure 3 displays 1731 links with small *p*-value (10^−700^), while the complete set of 17,873 links is listed in Supplemental Table S1. The communities analysis of the network, depicted in Fig. 3, grouped nodes by clumps of nodes that were more connected to each other than to the rest of the graph^21^; as result nodes were grouped in seven communities with acceptable value of modularity (*Q* = 0.49, Fig. 3). Next, we analyzed gene ontology terms (molecular function and biological process) of the genes associated to each community. In this manner, putative functions of uncharacterized genes can be inferred based on the genes with a known function in the same community.

**Figure 3.**
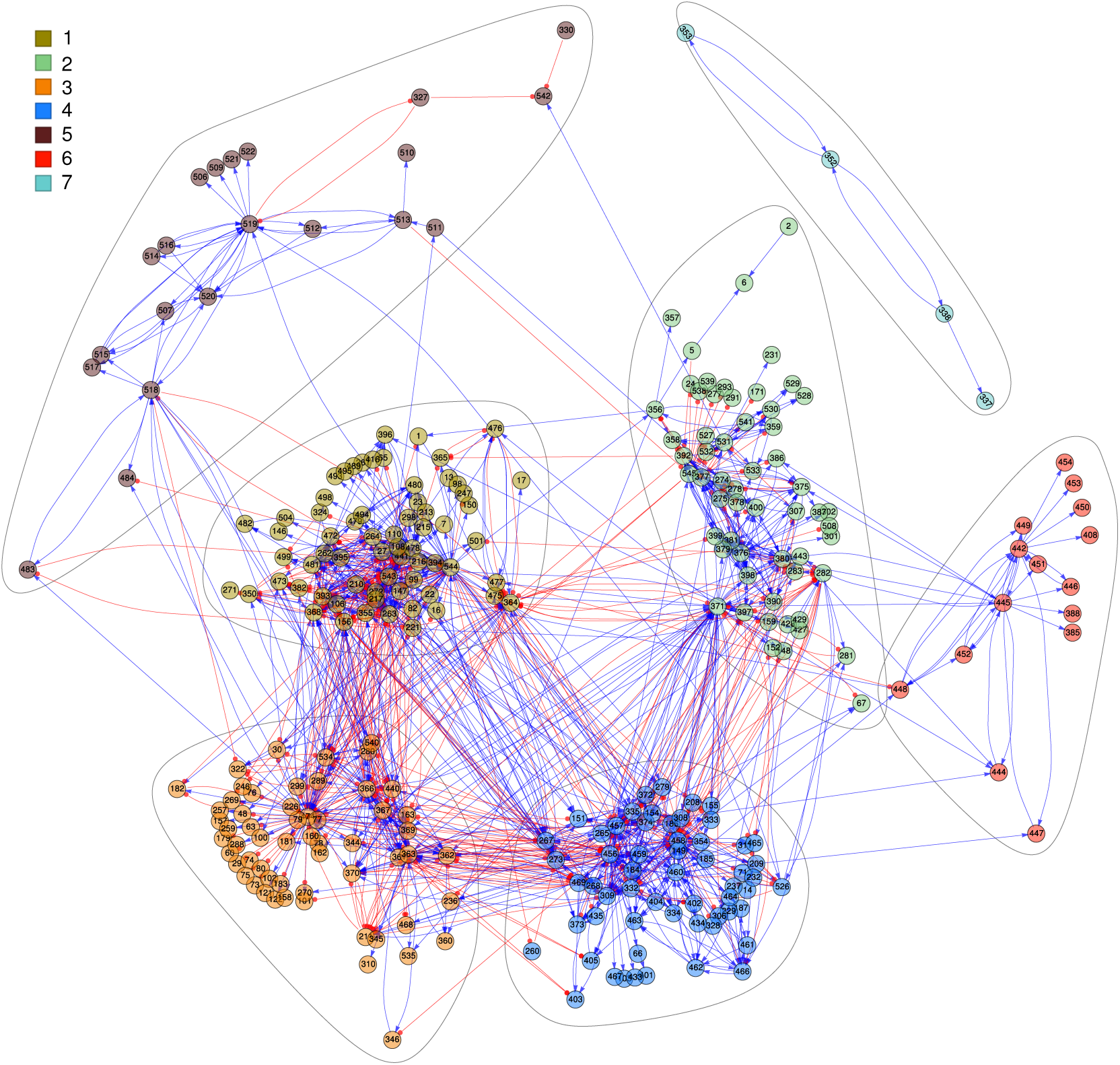
*T. gondii.* gene regulatory network. Cluster of genes (colored circles) are grouped in seven communities. By definition, nodes that are members of a community are more connected to each other than nodes belonging to other communities. Additional details of communities and nodes are in Supplemental Fig S1 and Supplemental Table S2.

Supplemental Fig S1 illustrates, by a word-cloud plot, the most representative terms for all communities, which are listed by extension in Supplemental Table S2. A total of 339 genes in the network are uncharacterized, whereby process and function can be assigned based on the nodes with known functions that integrate the community. Nodes in community 1 participate, mainly, in oxidation-reduction process and 44 uncharacterized genes are part of this community.

The analysis of community 2 indicates that 49 member genes, with no known function, could participate in the DNA repair process. Community 3, that contains 66 uncharacterized genes, is integrated of genes associated to a variety of processes, but translational elongation stands out. Community 4 genes are mainly associated to the biosynthesis of lipopolisaccharides, and include 50 uncharacterized genes. Community 5 contains 105 unknown genes while the rest of the genes are associated with proteolysis and cell adhesion. Interestingly, community 6 group genes participate in pathogenesis, include 25 yet uncharacterized genes. Members of this group should be studied in more detail since genes associated to this community could be may be novel pathogenic determinants which could give rise to the study of new pathogenic determinants. Finally, community 7 does not contain uncharacterized genes, but the genes product that integrate it participate in translation process. In conclusion, combining clustering methods and graph structure analysis it would be possible systematically assign processes and functions to a large group of genes in the network.

Extracting valuable information from a network made of more than 10,000 links can be a bottleneck in the genome-wide network analysis. One way to overcome this problem is by considering only the more important regulators of each steady state. Since the whole regulatory output of a gene depends on the gene’s activity level, some genes can be important regulators in one state, while their activity level in the other three states is low (i.e., *x*_*i*_ *∼*0). With this in mind, we have constructed network plots that emphasize the nodes with an important role as regulator in each steady state. In this sense we have display only those nodes that markedly regulate more than 5 other nodes. We have consider a regulatory interaction as marked when it explain more than 5% of the activity of regulated node, i.e., when *|w*_*i,* *j*_*x* _*j*_*| ≥* 5% of *|x*_*i*_ *|*. Thus, this feature depends not only on the weight of the link, but also on the current activity level of the regulator node. The Supplemental Fig S2-S6 depict the link-derived networks corresponding to parasite’s stages Od0, Od10, Tzd2, Bzd21 and Mc52, respectively. The main regulatory nodes represented in these plots were listed in Supplemental Table S3. We have found that the five analyzed stages share thirty-three of these nodes, twenty one of these nodes are present in the the network showed Fig. 3, and are listed in Table 1.

**Table 1.** Regulatory clusters common to all the parasite stages. The communities to which each cluster belongs and the most representative gene ontologies are detailed.

| Cluster ID | # genes | # marked links | Community | Molecular Function | Biological Process |
| --- | --- | --- | --- | --- | --- |
| 1 | 1 | 67 | 1 | GO:0003777 | GO:0055114 |
| 6 | 1 | 83 | 2 | GO:0006281 | GO:0006310 |
| 75 | 14 | 2 | 3 | GO:0003676 | GO:0006414 |
| 263 | 1 | 132 | 1 | GO:0003777 | GO:0055114 |
| 327 | 4 | 66 | 5 | GO:0004190 | GO:0006508 |
| 330 | 2 | 70 | 5 | GO:0004190 | GO:0006508 |
| 359 | 1 | 66 | 2 | GO:0006281 | GO:0006310 |
| 374 | 1 | 105 | 4 | GO:0004672 | GO:0006468 |
| 442 | 5 | 102 | 6 | GO:0004672 | GO:0009405 |
| 459 | 1 | 95 | 4 | GO:0004672 | GO:0006468 |
| 474 | 2 | 178 | 3 | GO:0003676 | GO:0006414 |
| 478 | 1 | 157 | 1 | GO:0003777 | GO:0055114 |
| 501 | 1 | 52 | 1 | GO:0003777 | GO:0055114 |
| 513 | 6 | 76 | 5 | GO:0004190 | GO:0006508 |
| 518 | 1 | 97 | 5 | GO:0004190 | GO:0006508 |
| 519 | 3 | 91 | 5 | GO:0004190 | GO:0006508 |
| 520 | 1 | 73 | 5 | GO:0004190 | GO:0006508 |
| 531 | 1 | 98 | 2 | GO:0006281 | GO:0006310 |
| 540 | 1 | 67 | 3 | GO:0003676 | GO:0006414 |
| 542 | 1 | 74 | 5 | GO:0004190 | GO:0006508 |
| 545 | 1 | 134 | 2 | GO:0006281 | GO:0006310 |

### Modeling the phenotypic transitions of *T. gondii*

The second step in our analysis is to include in the GRN dynamics the phenotypic transitions between the stages embedded in previous section. To this end we made a training set, denoted by *D*_*t*_, that considers the shortest possible path between the initial and final stages as indicated in Methods. For construction, the size of the training set is less than the size of GRN (*M < N*), consequently there are infinite solutions compatible with *D*_*t*_. Among them we have chosen a particular solution which is the closest one to the connectivity matrix **W**_*ss*_. Thus, the selected connectivity matrix, denoted by **W**_*t*_, can be found by:

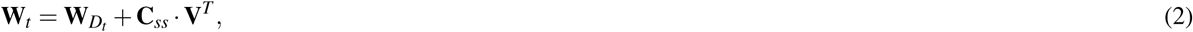

where **W**_*Dt*_ is the corresponding minimum *L*_2_-norm solution obtained by SVD for *D*_*t*_. Matrix **C**_*ss*_ was numerically computed by optimizing the overdetermined problem posed in the Method’ section (Eq.(10)). In this manner, the obtained connectivity matrix **W**_*t*_ is consistent with the phenotypic transitions included in training set *D*_*t*_, but also supports the multistability of the parasite’s life cycle. To test this ability we implement the model Eq.(1) with the connectivity matrix **W**_*t*_ to make simulations under different external cues. Each of these simulations was performed considering the associated external cue *µ* is acting. The simulations were performed by running 50 iterations of the GRN model initially in one of the parasite’s steady states, and storing the system’s state at each step. The temporal evolution of the 545 variables of the system can be illustrated by mean of the principal components or compiled in one movie. In Fig. 4 we plot 10 alternative trajectories for the phenotypic transitions Od0 →0d10, 0d10 →Tzd2 and Bz21 →Mc52, in the 3D space spanned by the main principal components. Each alternative trajectory, associated to the simulated phenotypic transition have the same initial condition, is affected by same external cue, but present a particular noise realizations. The final state of the system is in agreement with the expected state regarding the acting external cue. Alternatively, the complete temporal evolution of the system, from the oocyst to merozoite stage was compiled in a movie, which is available as Movie S1. Hence, the model can emulate the observed dynamical behavior of *T. gondii* during its life cycle.

**Figure 4.**
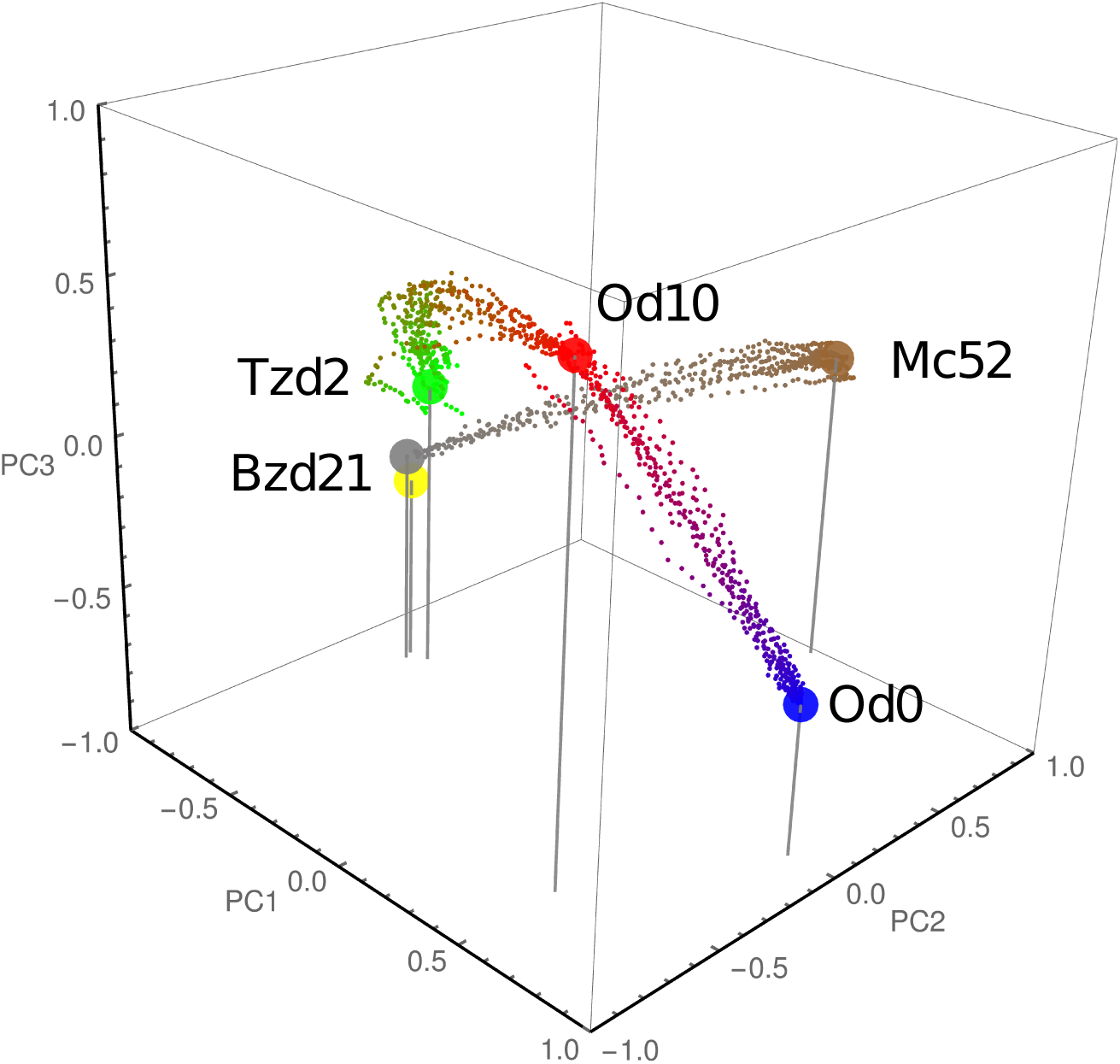
Phenotypic transitions between the steady states driven by external cues. The plot shows three phenotypic transitions of the system under the influence of external cues obtained with our model Eq.(1), in the space spanned by the three principal components. Each transition consist in 10 trajectories with different noise realizations and each trajectory has 44 intermediate states represented by small circles. Only the transitions Od0 → 0d10, 0d10 → Tzd2 and Bz21 → Mc52 were represented here. The transition Tzd2 → Bzd21 is better appreciated in Fig. 6A.

The phenotypic transitions are caused by an environmental signals, represented in our model by parameter **k**^*µ*^. To get insight about which genes could be modulated by external cues we have selected those clusters whose associate 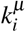-values are greater (lower) than the 95th (5th) percentile of the distribution. In this manner, we can identify the gene clusters which are activated (or inhibited) by the acting external cue *µ*. In this analysis we identify 140 gene clusters as externally regulated nodes, listed in Supplemental Table S4. This set comprises a total of 220 genes, 96 of which are still functionally uncharacterized. Interestingly, 40 of these genes are related with antigens, like microneme (MIC) proteins, dense granule (GRA) proteins, SAG-related sequence (SRS) proteins, and rhoptry proteins (ROP). Also two genes that coding for glycolitic enzyme enolases were indicated as externally regulated during transition Tzd2 →Bz21 by our analysis (clusters 364 and 376). It is important to note that enolases are recognized moonlight proteins, i.e., proteins that have dual functions^22, 23^; in *T. gondii* enolases fulfill a second function as transcriptional regulators implicated in parasite differentiation and cyst formation^24, 25^.

Further analysis was to identify the subnetwork that controls the parasite’s life cycle. This task involves the isolation of a small subset of nodes and regulatory links from a network of more than 17, 000 links. The number of possible subnetworks within a network of such a size is very huge and the evaluation of all subnetwork can result in an unfeasible task. In order to solve this problem, we reduced the search space. With this in mind, we considered only those circuits that involve nodes with many regulatory links. To that purpose we search for cyclic graphs, that containing only regulatory clusters, in matrix **W**_*t*_ and evaluate the ability of the module to emulate the parasite’s dynamics. In order to evaluate the dynamics of each module, we have considered a binary version of the model, where the variables *x*_*i*_ are binary and the system evolves following the equation:

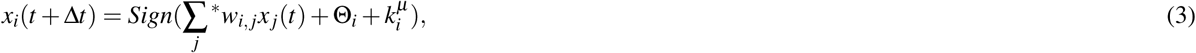

where * indicates that summation runs over the nodes belonging to the module. As final result we are able to recover a subnetwork with fifteen nodes which topology is illustrated in Fig. 5. The parameters values associated to this subnetwork are the same that the ones determined by Eq.(2), and they are listed in Supplemental Table S5. The above subnetwork was able to reproduces many features of the dynamics of *T. gondii* life cycle, such as the phenotypic transitions Od0 →Od10, 0d10 →Tzd2, Tzd2 →Bzd21 and Bzd21 →Mc52. The list of clusters that compose this subnetwork includes: 274, 327, 371, 442, 445, 460, 518 519, 520, 530, 531, 533, 540, 542 and 545, which in turn comprises a total of twenty six genes. Fifteen of these genes still have no assigned known function, while the rest of genes already characterized includes four genes associated with GRA proteins and other three genes associated with ribosomal proteins, and one coding a redoxin domain-containing protein. Additional information about these clusters is given in Supplemental Table S6.

**Figure 5.**
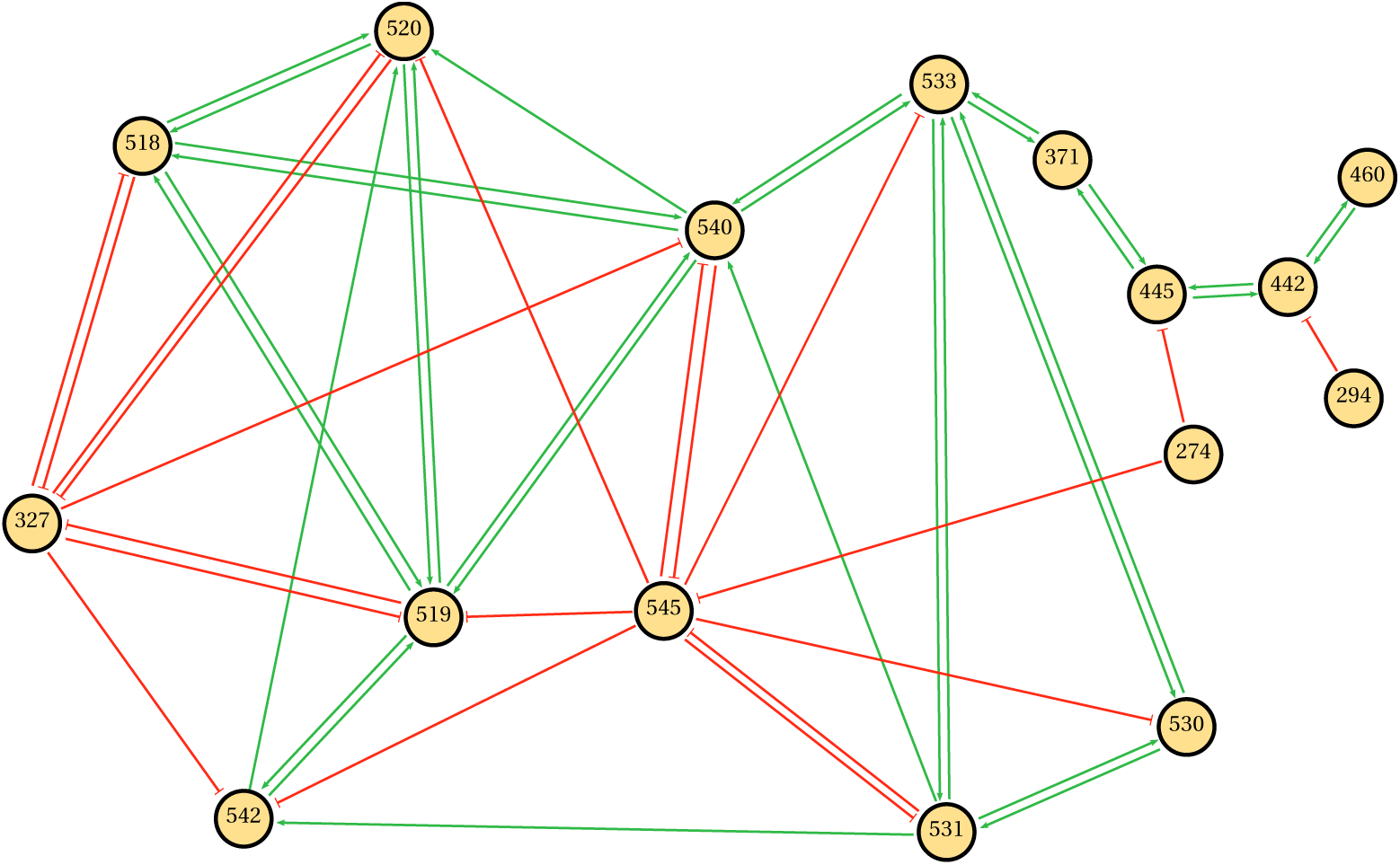
Architecture of the subnetwork linked to the parasite’s life cycle. This module is the minimal subnetwork that explain the studied phenotypic transition of *T. gondii*. Detailed information about node composition of this module can be found in Tables S8 and S9.

Finally, we perform *in silico* perturbations experiments on this module with the aim to confirm the relevant role of these nodes in networks dynamics. Since bradyzoite phenotype has an important role for the development of the chronic disease^2^, the perturbation analysis was focused over the tachyzoite to bradyzoite transition with the aim to identify relevant genes for the system’s dynamics. Table 2 summarize the results of the complete perturbation experiments over all subnetworks nodes in this transition. Figure 6 illustrates the influence of perturbation of node 274 in such phenotypic transition. While Fig. 6A depicts the transition of wild-type (WT) in the space spanned by the three principal components, Fig. 6B shows that deletion, or knock-out (KO), of 274 node prevents the system from reaching the bradyzoite stage. Additionally, the knock-down (KD) of this node drives the state of the system even far from the expected fate (Fig. 6C). Perturbations which are in the same direction of the WT does not impair to the system reach the expected fate. In this sense, Table 2 shown that there is always at least one perturbation without effect in th final fate, for example, the overexpression (OE) of the 274 node does not cause effect because this node is highly active during this transition. Other examples are shown in Supplemental Fig S7, where perturbation of clusters 519 (KO), 442 (KD) and 540 (OE) impairs the ability of the system to achieve the bradyzoite stage from tachyzoite. It should be noted that node 442 is composed mostly of genes that codify for dense granule antigens, see Supplemental Table S5. The above observation is interesting since these antigens are proposed as important factors to the development of the asexual phase of the cycle, particularly in tachyzoite and bradyzoite stages^26^.

**Table 2.** Summarize of *in silico* perturbation experiments over subnetwork nodes. The experiment was performed on the transition from tachyzoite to bradyzoite steady states. The cases in which the system reached the bradyzoite stage were indicated with the symbol “+” while those in which it was not reached were indicated with symbol “-”. KO: knock-out; OE: over-expression; KD: knock-down.

| Cluster ID | KO | OE | KD |
| --- | --- | --- | --- |
| 274 | - | - | + |
| 327 | + | - | - |
| 371 | + | + | - |
| 442 | + | + | - |
| 445 | + | - | - |
| 460 | + | - | - |
| 518 | + | - | + |
| 519 | - | - | + |
| 520 | + | - | + |
| 530 | + | - | + |
| 531 | + | - | + |
| 533 | + | - | + |
| 540 | + | - | + |
| 542 | - | - | + |
| 545 | + | + | - |

**Figure 6.**
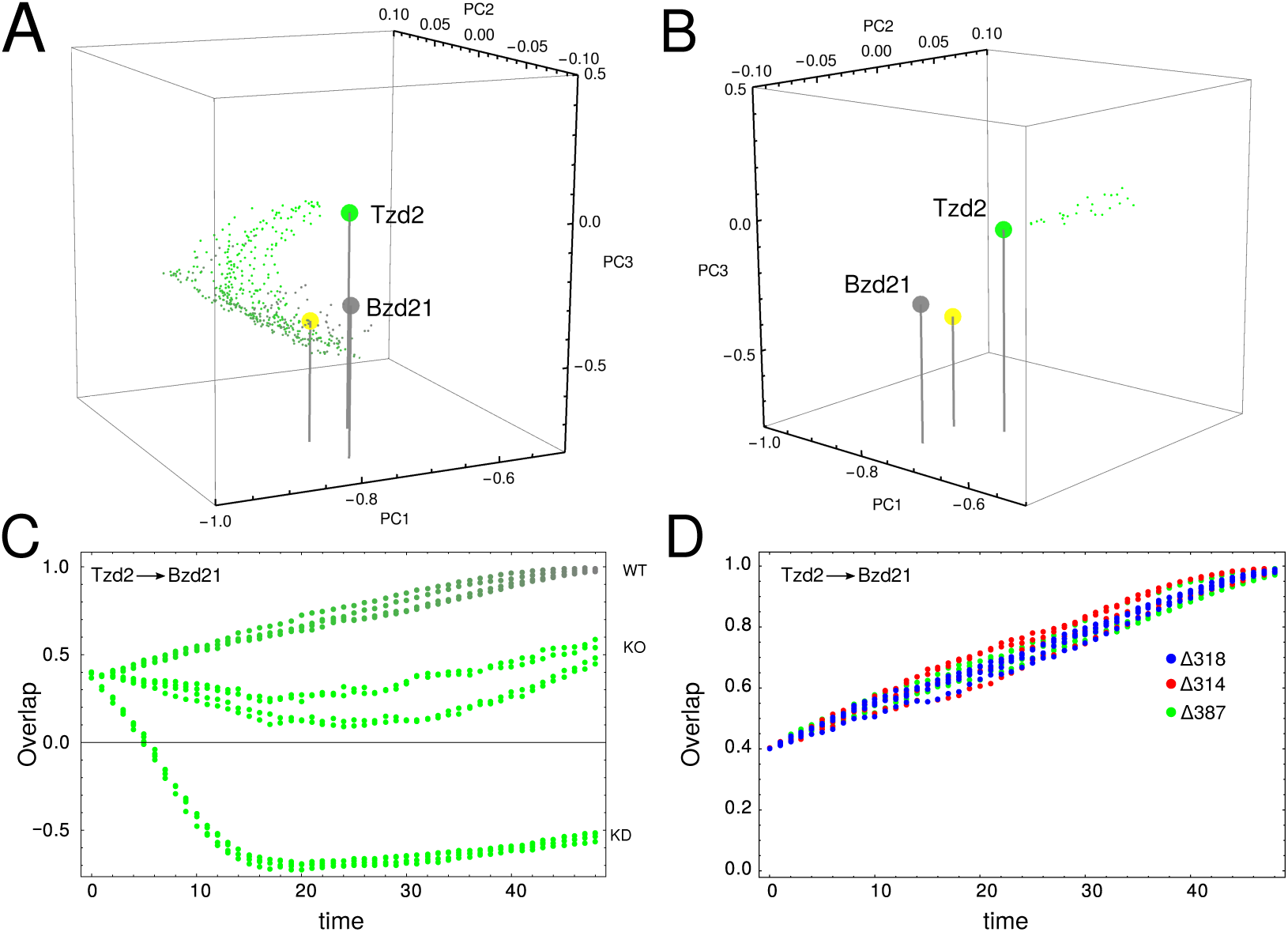
Phenotypic transition Tzd2→Bzd21. (A) A three principal components representation of Tzd2→Bzd21 transition obtained with our model Eq.(1). (B) Trajectory of the system when cluster 274 was deleted. Our model predicts that with this mutation that system can not complete the Tzd2→Bzd21 transition. (C) Temporal behavior of the overlap between the current state of the perturbed system and Bzd21 stage obtained with our model. We explore four alternative simulations of two different perturbations: knock-down (KD) and knock-out (KO). (D) Temporal behavior of the overlap between the state of the system at time *t* and the Bzd21 stage for our model with three different innocuous mutations.

In order to give significance to the above perturbation analysis of the module, we also perform perturbations on clusters which are not member of this subnetwork. In this sense we select at random thirty clusters, each one was analyzed by perturbation in terms of knock-out, overexpression and knock-down during Tzd2 →Bzd21 transition. Figure 6D shows three examples of these control perturbations, where the system reaches the bradyzoite stage. We found that in only 6 cases (*∼*6.6%) the perturbation have success into impair that system reach the final fate. Thus, the perturbation experiments suggest that the clusters proposed could be suitable candidates for master key regulators.

## Discussion

Protozoan pathogens like *Toxoplasma gondii* transit a complex life cycle’s during which the parasitic cell adapts to the context. Environmental signals of every particular context are interpreted by the cell through a intricate network of genes that allows the parasite cell to develop a phenotype for each context. Elucidating this network of genes would be relevant to understand the development of parasitic chronic diseases like Toxoplasmosis, contributing thus to the design of modern therapies and diagnostics for this affection.

In this work, we integrated microarray expression data from five different phenotypes of the life cycle of *T. gondii* in a GRN model. The information of the phenotypic transitions between the different stages was used to implement a reverse engineering procedure which allowed to reconstruct the connectivity matrix and determine parameter values linked to external modulations. From this matrix we were able to identified the key network module that drives the phenotypical transitions, as well as the genes target of the external modulations. In this way we embed the dynamics of the pathogen’s life cycle in a high-dimensional network system.

Clustering methods have been traditionally used to infer functions of uncharacterized genes^18^. Basically, genes with known function grouped with not yet characterized genes would allow inferring the functionality of these ones latter. In particular, experimental data have confirmed that genes participating in similar processes are co-expressed during the *T. gondii* replication cycle, even preserving the same *cys*-regulatory elements^27^. However, since of the 7,798 genes represented on the *T. gondii* chip 3,671 are not characterized, many clusters were completely integrated by uncharacterized genes. This feature, common in non-model organisms, can impair the gene functions prediction task, via clustering methods. Alternatively, in a previous study on the Apicomplexa *Plasmodium falciparum*, authors have assigned functions to thousand of uncharacterized gene modeling the parasite interactome, by using Bayesian approach^28^. Here we performed further analysis based on communities in the gene interaction network^21^ help to improve the gene annotation task. The identification of communities in a graph and the subsequent study of the structure of these communities allows to determine functional motifs within a molecular network^29, 30^. Our enrichment analysis over the biological process of every gene in each community reveals that particular process are predominant in different communities. By combining clustering and communities analysis could be possible infer the biological process of uncharacterized genes. Using this analysis we were able to predict the function of 339 uncharacterized genes. Thus, our extended gene clustering procedure could be useful to predict common *cys*-regulatory elements, design experiments for determination of protein-DNA interactions, and to improve our current knowledge of the transcriptional regulatory network, as in^13, 16^.

Furthermore, our framework was able to predict a key module that govern transitions between *T. gondii* steady states. This key network module is formed by twenty five clusters that could explain transitions between steady states. Most of the genes that integrate the master regulator of *T. gondii* are uncharacterized proteins. We have highlighted in this module the presence of dense granule proteins, as components of the cluster 442. GRA proteins constitute a group of relatively small proteins that are important to development and metabolism of parasitophorous vacuole, a highly dynamic compartment defining the replication permissive niche for the actively growing tachyzoite form of the parasite^26, 31, 32^. Our *in silico* perturbations experiments confirm that knock-out or overexpression of the cluster 442 do not prevent transition from tachyzoite steady state to bradyzoite steady state but knock-down of this genes could affect parasite cell fate when tachyzoite to bradyzoite transition is evaluated in our model. This observation is consistent with previous published results for GRA6 protein, were a biological role en cyst differentiation is proposed^33^. In addition, previous studies on a mutant Δ GRA2 strain are interesting since it tends to develop cysts *in vivo* unlike wild-type counterpart^34^. While that GRA1 its a essential factor to host invasion and replication^35, 36^. A similar perturbation analysis over thirty clusters selected at random, have scarce ability to alter the dynamics of the system, supporting the key regulatory role proposed by our modeling.

In conclusion, in this work we confirm that our previously mathematical approach its extrapolate to other protozoan pathogens allowing to reveals a subnet of master regulators that explain the dynamics of the life cycle transitions between phenotypes of *T. gondii*. Our results suggest that GRA proteins genes could act as key regulators in tachyzoite to bradyzoite differentiation. This finding is consistent with a previous study and reinforces the postulated role of GRA proteins in bradyzoite cysts development^37^. Consequently, experimental data based on perturbation experiments of the modeled network are necessary to confirm these observations. Finally, the methodologies employed here for the analysis of the modeled GRN could be useful to predict process and function of uncharacterized genes.

## Methods

### Data normalization

For this work we have used two microarray experiments made with the same chip^38^ and performed over type II clonal strains, M4 and TgNmBr1. In one of the studies Fritz and col.^39^ presents transcriptomics series of *in vitro* tachyzoite, *in vivo* and *in vitro* bradyzoite and complete oocyst development. On the other hand, in the second study Behnke *et al.*^40^ describes global gene expression of merozoite stage and integrate his results with the data obtained in the first study. Data sets are comparable^40^ and are publicly available in Gene Expression Omnibus (GEO) database (Accession no.: GSE32427 & GSE51780). This experimental series represents expression analysis of four of the five states that comprises the life cycle of T. gondii. Expression was evaluated per replicated at different times for each state and we selected the following data sets for analyze: oocysts day 0 (Od0) and oocysts day 10 (Od10, mature oocysts), tachyzoite day 2 (Tzd2), bradyzoite day 4 (Bzd4) and bradyzoite day 21 (Bzd21), and merozoite of cat #52 (Mc52). The chip used in both studies provide whole genome expression profiling, using at least 11 perfect match probes for each of the *∼*8000 genes in the *T.gondii* genome, including both the apicoplast and mitochondrial genomes. Also include a variety of controls (actin, hypoxanthine-xanthine-guanine phosphoribosyl transferase, yeast housekeeping genes and mismatch probes), immune effector molecules (cytokines, receptors, etc), and genes whose expression is suspected from previous to be altered by infection. Further details about probe preparation, microarray hybridization, and data acquisition can be found in^38^, while the description of the microarrays is available at http://ancillary.toxodb.org/docs/Array-Tutorial.html. Microarray data of the two data sets were loaded into R software using the *affy* package from Bioconductor Project and processed using Robust Multi-array Average (RMA) and quantile normalization^41^. The signal relative intensity of the probe *i* recorded in one of the biological replicates, *j* = 1, 2; at one of the time point *α* = 1, 2, 3, 4, 5, 6; was represented by 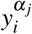. After within-slide replicates processing, we averaged the relative intensity 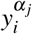. over all replicates belonging to the same time point (stage), i.e 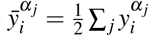 Control data was eliminated and only expression data from specific *T. gondii* probes were analyzed, which give us a normalized expression set of 7, 798 probes for each sample. Thus, we considered the variable that describes the expression level of the probe *i* at time point *α* as the quantity 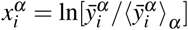. supplemental Table S7 lists all normalized expression levels, 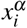, used in the next step, with their corresponding probes identifiers (probe ID).

### Redundancy reduction procedure

With the aim of group the genes by similar expression profiles, we have applied an agglomerative hierarchical clustering method known as unweighted pair group method with arithmetic mean (UPGMA). The agglomerative process is stopped at a given number of clusters, *N*_*c*_, considered suitable for our data-set^42^. This suitable number of clusters is not known beforehand, and it needs to be estimated. In order to do this, we repeated the clustering procedure for several *N*_*c*_ values, and computed the measure of the clustering merit, known as Davies-Bouldin index (DBI)^43^. This index is defined as:

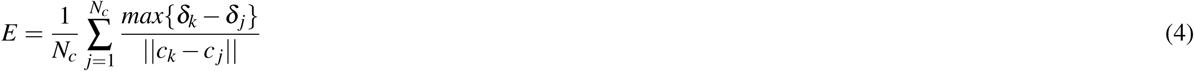

where ‖*c*_*k*_ *- c* _*j*_‖ is the distance between the cluster centroids, and 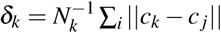 denotes the centroid intra-cluster distances of cluster *k* (*N*_*k*_ being the number of genes belonging to cluster *k*). Low DBI-values indicate good cluster structures. It is true, of course, that increasing *N*_*c*_ without penalty will always reduce the resulting DBI-values. Then, in the choice of a suitable *N*_*c*_ the trade-off usually involves a balance between the data compression and the accuracy of the dimensionality reduction. Supplemental Fig S8 displays the DBI as a function of *N*_*c*_ for the gene-expression profile under study. It can be seen that the DBI does not suffer a significant reduction beyond *N*_*c*_ = 545. Thus, we selected this value as the optimal number of clusters. The profiles of the 7, 798 genes were grouped in 545 clusters, and the intra-cluster average of the expression level (i.e., 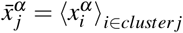) was used as dynamical variable for the subsequent modeling. The cluster membership of each genes is listed in Supplemental Table S8, while the intra-cluster averages of the expression levels, 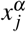 for each cluster are listed in Supplemental Table S9 for each time point. The resulting average levels, 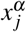, are illustrated in a 2D array plot for the six stages of *T. gondii* life cycle in Fig. 1B.

## Reverse engineering methods

### The gene network model

Linear models serve as the basis of all continuous gene-network approaches currently available to model typical time-course gene-expression data sets^18, 44–46^. In this work we use a discrete-time version of this linear network model which owns two interesting advantages: its parameter estimation does not involve intensive computational steps^18^, and it can take into account fluctuations. In this network model, the state of the system at time *t* is determined by the activity of the *N* nodes of the network, represented by a *N*-dimensional vector **x**(*t*). The equation governing the temporal evolution of the linear GRN can be written as:

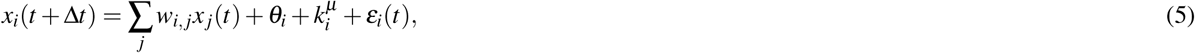

where we have added a noise term, *ε*(*t*), assumed to be Gaussian with mean equal to 0. *w*_*i,* *j*_ are the elements of the weighted connectivity matrix **W**, *θ*_*i*_ is a constant bias term to capture the activity level of gene *i* in the absence of regulatory inputs, and 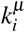 determines the influence of the environmental cue *µ* over gene *i*. Notice that the bias term and the acting environmental cue can be included in an extended version of matrix **W** and of state vector *x*. Thus, in order to simplify the notation, the state of gene *i* can be written as:

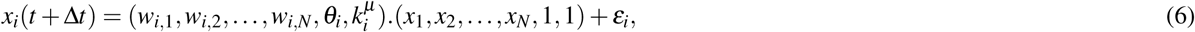

where *µ* corresponds to the acting environmental cue. This said, the same parameter estimation method can be applied whether the environmental cues are present or not.

### Parameter estimation

In order to estimate the parameter values associated to *w*_*i,* *j*_, *θ*_*i*_ and 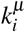 we give in this section some definitions. We assume that our available gene-expression data consist of *M* pairs of input-output states, represented by *D* = {**X**, **Y**} The input matrix **X** is the *N ×M* gene-expression matrix, which columns **x**_*v*_, correspond to the input states at time *t*, while the rows indicate individual genes. The same is valid for the gene-expression matrix at time *t* + Δ*t*, **Y**. For a given *D*, the linear model must map each gene-expression state to the consecutive state, i.e.:

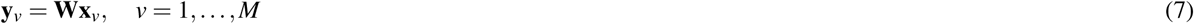

In order to find the matrix **W** we minimize the cost function 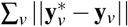, in such a way that the predicted state, 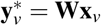 must be as close as possible to the associated output state **y**_*v*_. This minimization problem has many alternative solutions when *M < N*, one of them is the one with the smallest *L*_2_ norm, denoted here by **W**_*L*2_. This solution is given in terms of the singular value decomposition (SVD) of matrix *X*, i.e. **X**^*T*^ = **U S V**^*T*^, where **U** is a unitary *M × N* matrix of left eigenvectors, **S** is a diagonal *N ×N* matrix containing the eigenvalues *s*_1_, *…, s*_*N*_, and **V** is a unitary *N ×N* matrix of right eigenvectors^45, 47^. Of course the superscript ^*T*^ denotes the transpose matrix. Thus, the minimum *L*_2_ norm solution *W*_*L*2_ is given by:

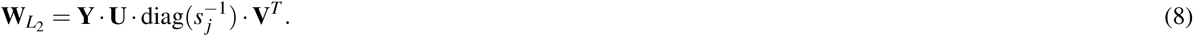

Assuming that **x**^*v*^ are linearly independent, finding the unique solution requires that *M ≥N*. However, the inverse problem in GRN involves *M ≪ N*. Thus, the problem tends to be severely undetermined, and many solutions are consistent with the data set *D*. Fortunately, there exist a closed-form expression for all possible connectivity matrices which are consistent with Eq.(7) in terms of **W**_*L*2_:

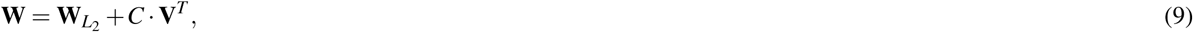

where **C** is an *N × N* matrix whose elements *c*_*i* *j*_ are 0 as long as *s* ≠ _*j*_0. Otherwise, they are arbitrary scalar coefficients. As it will be seen later, the degrees of freedom due to this arbitrariness can be exploited to our benefit^12^, 45. In the first step, the solution offered by Eq.(8) will be implemented to embed the six stages of the life cycle *T. gondii* into the dynamics of the GRN model, without considering the transitions between these states. In a second step, Eq.(9) will be used to uncover the environmental cues by means of using the information provided by the phenotypic transitions, and the connectivity matrix associated with the steady states inferred in the first step.

### Embedding the steady states

In this first step of the inferring procedure we construct a training set *D*_*ss*_ with size *M*. To this end different noise realizations associated with each stage *α* were added to obtain the columns of matrix **X** as follow:

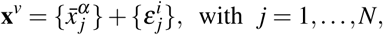

where the index *α* runs from 1 to 6, the index *v* runs from 1 to M, and the superscript *i* denotes the noise realizations. *ε* _*j*_ is a Gaussian noise with mean equal to 0 and a small standard deviation (set at 1% of the expression data). In this work, we have used 50 noise realizations for each state. Thus, *M* = 6 *×* 50 = 300. The columns of matrix **Y** are defined in the same way, i.e 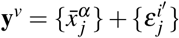, but with different noise realizations, of course. Thus, we have expanded the training set size, thereby making the steady states more robust against the fluctuations. This implies that if at a given time the system is close to one steady state, it will remain close to that steady state in the next time-step as well. This procedure allows the model to display the same set of steady states as the ones seen in the parasite.

In order to obtain a sparse connectivity matrix we need to discriminate whether the estimated matrix elements *w*_*i,* *j*_ are 0 or a value significantly different from 0. To this end, we performed a location test for each *P*(*w*_*i,* *j*_) distribution, as described in^12^. Basically, we construct an ensemble of training sets and for each set we have an associated slightly different solution. This ensemble of solutions defines a probability distribution for each weight, *P*(*w*_*i,* *j*_). If the *p*-value associated with the location test is greater than 0.01, the hypothesis is accepted and the mean value of *P*(*w*_*i,* *j*_) is assigned to *w*_*i,* *j*_, otherwise, *w*_*i,* *j*_ = 0. This procedure allows us to obtain a sparse connectivity matrix, compatible with the steady states, which will be denoted by **W**_*ss*_.

### Embedding the phenotypic transitions

In the second step, we extend our analysis to embed the transitions between the states and determine the environmental cues which drives the transitions. In this sense we used the extended versions of **W** and **x** described by Eq.6. We assume that these transitions take place gradually through the shortest path between the consider two stages *α* and *β*. In this manner, if the state of the system is driven from state *x*^*α*^ to the state *x*^*β*^ due to an external signal *µ* = *β*, then the system makes a series of small transitions between intermediate states denoted by *x*^*α,β*^ (*t*). We construct these intermediate states by means of a linear combination of the initial and final states as follow:

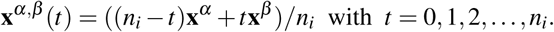

By using these intermediate states, we make a new training set for embed the transitions *α →β*, denoted by *D*_*t*_. Thus, the matrix’ columns **x**^*v*^ and **y**^*v*^ are given by:

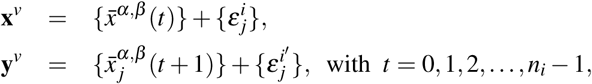

where *ε* _*j*_ is a Gaussian noise with mean equal to 0 and a small standard deviation (set at 1% of the expression data). In this paper we have consider four phenotypic transitions: Od0 → Od10, Od10 → Tzd2, Tzd2 → Bzd21, and Bzd21 → Mc52. The transition Tzd2 → Bzd21 includes the stage Bzd4 as part of the transition trajectory. For each transition, we consider 44 small transitions, i.e. *n*_*i*_ = 44. Consequently, the size of training set *D*_*t*_ was *M* = 176, since *M < N* there exist many solution consistent with *D*_*t*_. However, we are interested in the particular solution which as close as possible to the previously determined **W**_*ss*_. In order to determine this solution, we computed the smallest *L*_2_ norm solution associated to *D*_*t*_, denoted by **W**_*Dt*_, and by using Eq.(9) we estimate the matrix **C**_*ss*_ by mean of the equation:

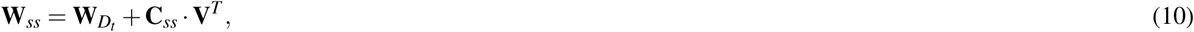

To determine the elements of matrix **C**_*ss*_ from Eq.(10) is an overdetermined problem. This kind of optimization problem can be addressed by applying the interior point method for *L*_1_ regression^45^. The resulting *c*-values were then used to compute the particular solution, represented by **W**_*t*_, by using **W**_*t*_ = **W**_*Dt*_ + **C**_*ss*_ **V**^*T*^. This new connectivity matrix is consistent with the phenotypic transitions included in *D*_*t*_, but it is also very close to **W**_*ss*_.

## Acknowledgements

A.M.A. is a postdoctoral fellow of the CONICET (Argentina). M.M.C. and L.D. are researcher members of the CONICET (Argentina). L.D. acknowledges financial support from Consejo Nacional de Investigaciones Cientificas y Tecnicas, Project No. PIP 0597 (Argentina).

## Author contributions statement

A.M.A. and L.D. conceived and performed experiments. M.M.C and L.D. analyzed the results and wrote the manuscript. All authors reviewed the manuscript.

## Legends of Supplementary Tables

**Supplemental Table S1:** Regulatory links. List of the 17,873 significant weights (at a significance level of 0.01) needed for the maintenance of the parasite’s steady states. The first column indicates the regulatory cluster IDs. The second column indicates the regulated cluster IDs. The third column lists the mean values of each cluster, averaged over the ensemble of 500 training sets. The fourth column lists the associated standard deviations. The last column lists the *p*-values (the probabilities of the location test).

**Supplemental Table S2:** List of genes grouped by related functions in 7 communities.

**Supplemental Table S3:** Main regulatory genes. List of the genes with important regulatory activity for the maintenance of the parasite’s steady states.

**Supplemental Table S4:** Main regulated genes. List of the genes regulated by the external cues responsible for the transitions between the steady states.

**Supplemental Table S5:** Regulatory links of *T.gondii*’s life cycle subnetwork. Estimated values of parameters *w*_*i,* *j*_, Θ_*i*_, and 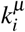, extracted from matrix *W*_*t*_.

**Supplemental Table S6:** Master key regulators. List of genes that integrate the clusters of the subnetwork that regulate the *T. gondii*’s life cycle.

**Supplemental Table S7:** Gene-expression profile of *T. gondii*’s life cycle. Log-norm expression levels corresponding to 7,798 *T. gondii* genes, obtained from microarray experiments as indicated in Methods. The gene IDs listed in the first column correspond to our own gene numbering. The second column lists the microarray probe IDs. The third and the fourth columns list the *Toxoplasma gondii* data base (www.toxodb.org) IDs and protein names, respectively. The last four columns correspond to the gene-expression levels in each stage of the parasite’s life cycle: oocyst d0, oocyst d10, tachyzoite d2, bradyzoite d4, bradyzoite d21, merozoite, respectively.

**Supplemental Table S8:** Cluster composition. Clusters (cluster ID) are listed in the first column, with their corresponding genes (gene ID) listed in the second column. The third and the fourth columns list the gene and protein names, respectively.

**Table Supplemental S9:** Intra-cluster averages of the expression levels. Gene-expression levels of each cluster (rows) in each different stage (columns) used in all further modeling computations.

